# End-of-outbreak determination under seasonal transmission

**DOI:** 10.64898/2026.08.03.742445

**Authors:** WS Hart, C Mills, M Manica, F Menegale, M Spaziante, F Vairo, P Poletti, G Guzzetta, RN Thompson

## Abstract

At the apparent end of an infectious disease outbreak, policymakers must decide when to relax interventions, balancing epidemiological safety and economic efficiency. Here, we extend a renewal equation framework for estimating the risk of additional cases to scenarios with seasonal transmission, such as *Aedes*-borne virus transmission in temperate settings. In simulations, we demonstrate that the time of year is key to when it is safe to relax interventions under seasonal transmission: measures can be relaxed sooner if the last observed case occurs towards the end of the transmission season. We apply our framework to the 2017 chikungunya outbreak in Italy, integrating incidence data with estimates of temperature suitability for transmission, and further extend our approach to account for case under-reporting. Our model incorporating an end-of-season decline in transmission indicates a lower risk of additional cases late in the outbreak than a model based on incidence alone: assuming 60% case reporting, the seasonal model reaches a 1% risk threshold 22 days after the last observed case, compared with 39 days for the incidence-only model. Incorporating seasonality in model-based assessments of transmission risks at the end of an outbreak can therefore support more timely and reliable decision making.

## Introduction

Climate change is contributing to the ongoing expansion of the geographic range of *Aedes albopictus* and *Aedes aegypti*, which are vectors of the pathogens that cause diseases including dengue and chikungunya [1]. As a result, previously unaffected locations have begun to experience outbreaks of these diseases [2]. For example, chikungunya outbreaks driven by *Ae. albopictus* occurred in Italy in 2007, 2017 and 2025 [3–5]. In temperate settings, the potential for *Aedes*-borne pathogen transmission is seasonal, with summer conditions supporting mosquito activity and viral replication, but cold winter temperatures generally precluding sustained transmission [6,7].

Towards the end of an infectious disease outbreak, a key consideration for policymakers is when it is safe to relax or remove interventions. During *Aedes*-borne viral outbreaks, public health measures may include vector and clinical surveillance, vector control, public health messaging promoting personal protective measures, suspension of blood donations, and screening tests on available blood supplies implemented to prevent transmission via transfusions [8,9]. Maintaining interventions is often costly [10], but relaxing measures too early leads to an unacceptably high risk of further transmission occurring. Timely relaxation of interventions following the end of an outbreak is therefore essential.

For a range of diseases, including both vector-borne and directly transmitted diseases, the decision to relax interventions is typically based on a fixed waiting period following the last observed case [11]. For example, for *Aedes*-borne disease outbreaks, the European Centre for Disease Prevention and Control (ECDC) recommends a 45-day waiting period before the assigned risk level is lowered, at which point interventions can be relaxed [8]. This 45-day period comprises a theoretical maximum possible serial interval of 41 days, plus a four-day buffer for case diagnosis [8]. Similarly, for Ebola virus disease, the World Health Organization recommends that an outbreak can be declared over and interventions relaxed 42 days after the resolution of the last observed case [12]. However, modelling studies have shown that the risk of additional cases depends on outbreak-specific factors, including the time series of recent disease incidence [13], the time-dependent reproduction number [14] and the extent of case under-reporting [15]. Consequently, quantitative methods have been proposed to inform end-of-outbreak decision making by estimating the risk of additional cases: the probability that cases arise on or after a given day, given the disease incidence time series up to that day [11,13–30]. The risk can be recalculated each day or week, so that policymakers can relax interventions once the risk falls below a threshold chosen to balance the residual risk against the benefit of relaxing interventions sooner.

Here, we extend model-based approaches for estimating the risk of additional cases to settings with seasonal transmission, focussing on *Aedes*-borne disease outbreaks in temperate settings. Previous studies have examined environmental seasonality in the context of the risk that pathogen introductions lead to a large outbreak [31–34], highlighting the importance of accounting not only for transmission conditions at the time of introduction, but also for subsequent changes in transmission. Specifically, estimates based only on conditions at the time of introduction can underestimate *Aedes*-borne disease outbreak risks in the spring (as transmissibility increases following introduction) and overestimate them in the autumn [31,33,35]. While similar principles are likely to apply to estimates of the risk of additional cases towards the end of an outbreak, this has not to our knowledge been explored previously.

To address this, we extend a previous renewal equation-based approach for estimating the risk of additional cases [14] to incorporate seasonal changes in transmission. Using simulations, we show that in the presence of seasonality, the time of year of the last observed case is a key determinant of when interventions can be relaxed. We then apply our approach to chikungunya outbreak data from the town of Anzio, the epicentre of the 2017 Italy outbreak [36]. We compare risk estimates based on time-dependent reproduction numbers inferred from incidence data alone with those additionally informed by seasonal temperature variation, initially assuming full case reporting before generalising our approach to incorporate case under-reporting. Since the last case in Anzio was observed towards the end of the potential transmission season, neglecting the successive decline in transmission following the last observed case would lead to overestimation of the risk of additional cases and therefore interventions potentially being maintained longer than necessary. These findings highlight that considering seasonal variations in transmission in data-driven epidemiological modelling analyses can support timely public health decision making during outbreak surveillance and response.

## Results

Our approach combines three inputs: the disease incidence time series up to the current day (Figure 1A), the time-dependent reproduction number (*R_t_*, which captures seasonal changes in transmission and can be informed by historical data; Figure 1C) from that day onwards, and the serial interval distribution (governing the time between symptom onsets in infector-infectee transmission pairs; Figure 1B). From these inputs, we calculate the risk of additional cases, i.e., the probability of at least one further case occurring from that day onwards (assuming no additional imported cases; Figure 1D). Under the assumption that cases arise according to a discrete-time stochastic renewal equation model, an analytical expression for the risk can be derived. As we demonstrate, in scenarios where *R_t_* is inferred using Bayesian inference, uncertainty in *R_t_* can be incorporated into calculations of the risk of additional cases. For details of our approach, see Methods. Estimates of the risk of additional cases can then be used to inform the timing of decisions to relax interventions (Figure 1D).

**Figure 1.**
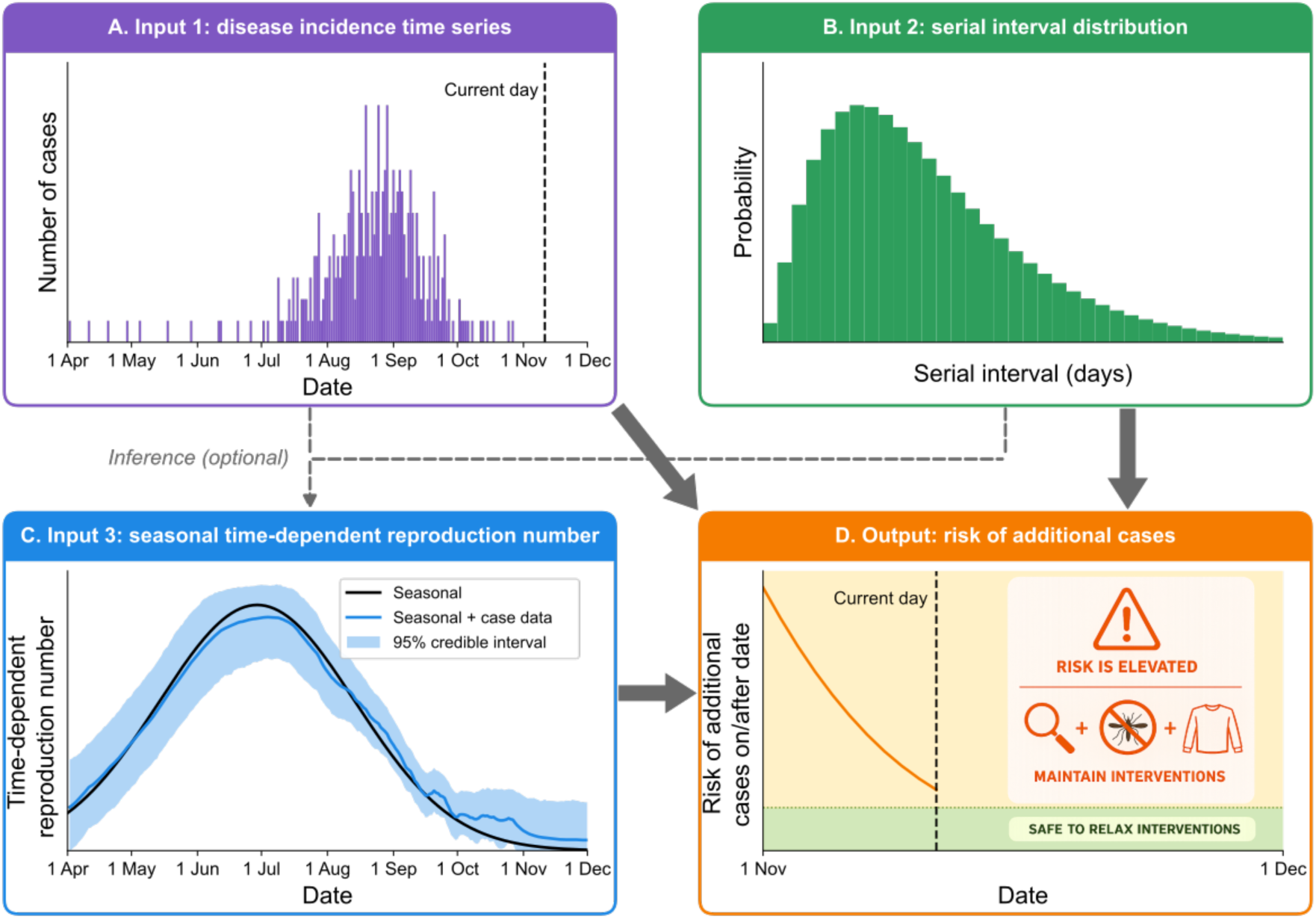
Schematic illustrating our framework for estimating the risk of additional cases towards the end of a seasonal outbreak. A-C. Inputs to our framework. A. The disease incidence time series up to the current day. B. The probability mass function of the serial interval. C. The time-dependent reproduction number, *R_t_*, which captures seasonal forcing (black curve) and may also be informed by inputs A-B using Bayesian inference (blue curve and shaded area). D. From these inputs, we compute, using the disease incidence data available up to a given day, the probability of at least one additional case on or after that day. This output can inform when interventions can be relaxed. For example, interventions could be maintained until the risk of additional cases falls below a specified threshold.

### Simulation study

To demonstrate that the time of year can be an important factor in decision making towards the end of an outbreak in the presence of seasonality, we initially considered a theoretical scenario in which *R_t_* varies seasonally (Figure 2A). Specifically, this seasonal transmission profile was chosen to be illustrative of *Aedes*-borne disease transmission in temperate settings, with sustained transmission only possible in warmer months (see the “Simulation study” section of the methods).

**Figure 2.**
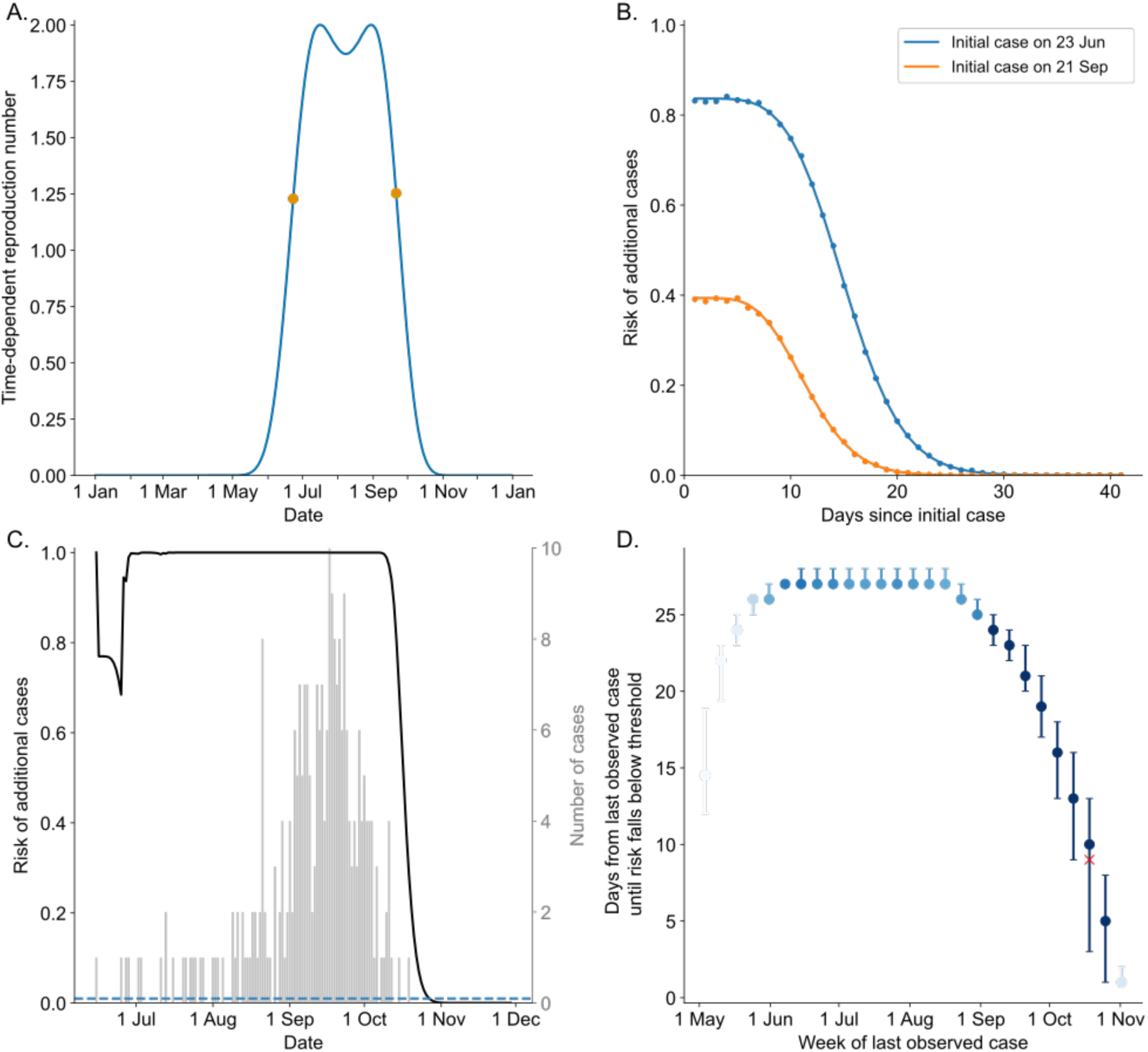
Simulation study of end-of-outbreak decision making for seasonal pathogens. A. Assumed seasonal profile of *R_t_*. Orange dots show the similar values of *R_t_* on 23 June and 21 September. B. Risk of additional cases following a single imported case on either 23 June (blue) or 21 September (orange), calculated either analytically (our default approach; lines) or via renewal equation model simulations (dots). C. Daily risk of additional cases for a simulated outbreak based on disease incidence observed prior to the date shown on the x-axis (bars show simulated incidence data). The horizontal dashed line indicates a probability of 0.01. D. Number of days from the last observed case to the risk of additional cases falling below 1%, shown by week of last observed case across 100,000 simulated outbreaks. Each outbreak was initiated with a single incident case on a day of year chosen uniformly at random, retaining only outbreaks in which at least one additional case occurred. Dots and whiskers show the mean and central 95% prediction interval, respectively, with colours indicating the proportion of simulated outbreaks in which the last observed case occurred in different weeks (darker corresponds to a higher proportion, with the colour saturating at 1% of simulated outbreaks). The red cross represents the example outbreak shown in panel C.

First, we compared two pathogen introduction times on either side of the assumed summer transmission peak (orange dots in Figure 2A), calculating, for each day following the initial case, the risk of additional cases on or after that day (assuming no cases arose in the intervening period, and assuming the *R_t_* curve to be known exactly; Figure 2B). Although *R_t_* is almost identical on the two introduction dates, we found a much higher risk of additional cases following the earlier introduction date, due to the increase in *R_t_* after the earlier date, whereas *R_t_* declines after the later date. This highlights that the precise timing of an outbreak in the year can substantially influence the risk of additional cases.

Then, to explore the consequences of seasonality for end-of-outbreak decision making in more detail, we used the renewal equation model to generate a large number of synthetic outbreaks (a single example outbreak is shown in Figure 2C). Each outbreak was initiated with a single imported case, introduced on a randomly selected day of the year but with only introductions resulting in local transmission being retained. For each outbreak, we calculated the number of days from the last observed case until the risk of additional cases fell below a 1% threshold. In principle, such a risk level criterion could be used to guide a decision to relax interventions. While we considered a 1% threshold as an illustrative example, policymakers could choose an alternative threshold according to their level of risk aversion. Figure 2D shows the dependence of the waiting time for the risk to fall below the 1% threshold on the time of year of the last observed case. We found that this waiting time is relatively consistent over simulations with a last observed case during the summer, but then rapidly declines when the last observed case occurs later into the autumn.

We note that, between outbreaks in which the last observed case occurs in the same week, waiting times are more variable in mid autumn than in summer (Figure 2D). This is because outbreaks ending in summer are typically small (since whilst *R_t_* is substantially above one, stochastic extinction is unlikely for all but very small outbreaks), whereas outbreaks of a range of sizes end in mid autumn, and the risk of additional cases occurring depends on the number of recent cases that have occurred.

To explore the dependence of our findings on the precise seasonal transmissibility profile, we conducted sensitivity analyses in which we varied either the maximum annual *R_t_* value (Figure S1A- C) or the rate at which transmission declines away from the middle of the summer transmission season (Figure S1D-F). If transmission remains possible later into the autumn and early winter, the waiting period until the risk falls below the 1% threshold is longer when the last observed case occurs in the autumn (Figure S1E). Nonetheless, in every scenario considered, the time of year of the last observed case strongly influences when model-informed decisions to relax interventions would be made.

### Anzio chikungunya outbreak case study

We then applied our approach for computing the risk of additional cases to data from the 2017 chikungunya outbreak in Anzio, Italy. This municipality was the epicentre of an outbreak that included multiple smaller clusters in the Lazio region [37], where Anzio is located, with a further large outbreak occurring in Guardavalle Marina in the Calabria region, over 600 km to the south.

To inform our framework with estimates of seasonality in transmission, we combined recorded temperature data (Figure 3A) with an existing model [38] describing temperature suitability for chikungunya transmission by *Ae. albopictus* (i.e., the temperature dependence of the relative reproduction number, up to a scaling factor capturing non-temperature determinants of transmission; Figure 3B), obtaining a curve describing how temperature suitability varies seasonally (Figure 3C). We used a smoothed temperature curve fitted to data from multiple years, since (i) our framework requires information about how transmission conditions are likely to change in future (i.e., complete data for the current year would not be available if applying our approach in real time), and (ii) vectors may not respond immediately to day-to-day variation in temperatures about a seasonal trend; however, as described below, we accounted for deviations in suitability from the seasonal trend curve during the outbreak based on transmission observed in the disease incidence time series.

**Figure 3.**
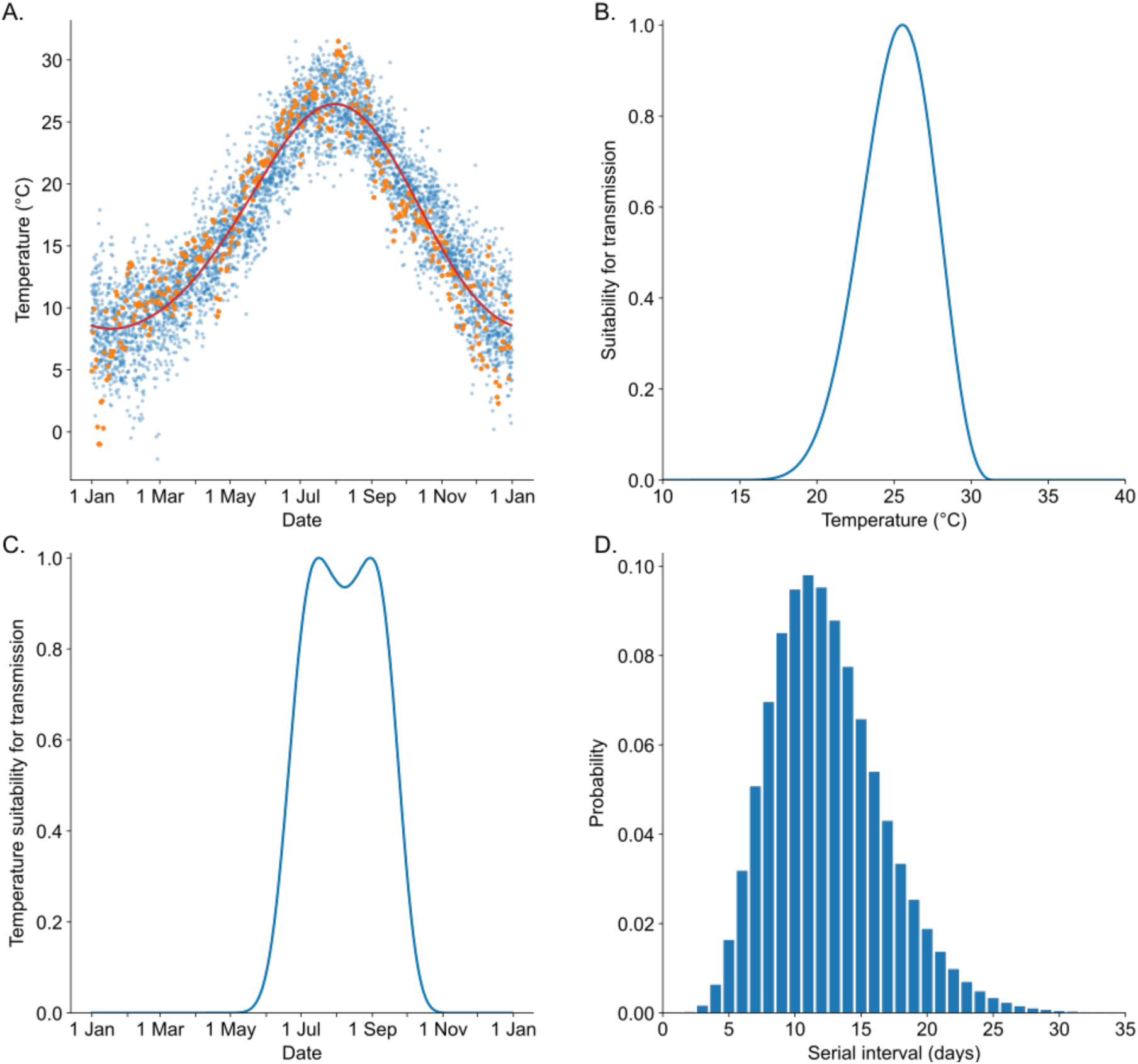
Inputs to our framework for estimating the risk of additional cases during the 2017 chikungunya virus outbreak in Anzio, Italy. A. Smooth periodic curve (two-harmonic Fourier model; red curve) fitted to daily temperature data for 2010-2024 (dots; values for 2017 shown in orange and other years in blue). B. Suitability for chikungunya virus transmission by *Ae. albopictus* at different temperature values (i.e., temperature-dependent components of the reproduction number, scaled to have maximum value one), as estimated by Tegar *et al.* [38]. C. Temperature suitability for transmission on each day of the year, based on the smooth temperature curve in A. D. Assumed probability mass function of the chikungunya serial interval (comprising the entire host-vector-host transmission cycle) [37].

We then fitted the renewal equation model to the Anzio disease incidence data (grey bars in Figure 4A-C). Specifically, we assumed a time-dependent reproduction number of form *R_t_* = *C_t_S_t_*, where *S_t_* describes temperature suitability for transmission and the scaling factor *C_t_* captures non- temperature determinants of transmission (for example, interventions such as vector control). We estimated both *S_t_* (Figure 4A) and *C_t_* (Figure S2). Estimates of *S_t_* were informed by the seasonal trend curve in Figure 3C and largely follow this trend for most of the outbreak (Figure 4A). However, the estimated suitability diverges from the trend curve around October 2017, with a positive value up to and beyond the last observed case on 5 November (despite the seasonal prior value being very small by this time) reflecting that transmission continued to take place when this would be unlikely based on the trend curve alone.

**Figure 4.**
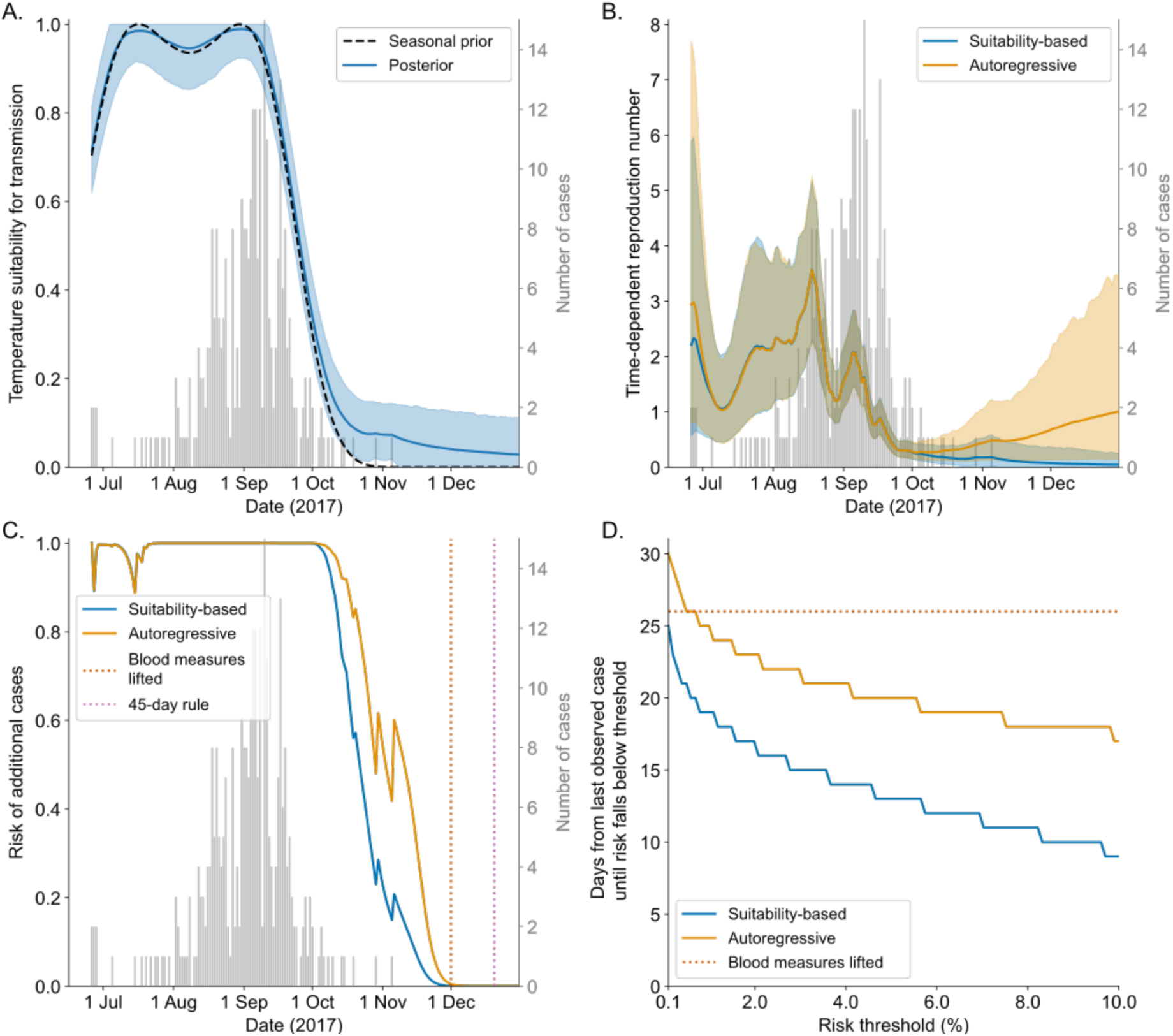
Application to the 2017 chikungunya outbreak in Anzio, Italy. A. Temperature suitability for transmission, *S_t_*, assumed a priori to follow an autoregressive process of order 1 (AR(1) process) centred on the seasonal profile estimated in Figure 3C (shown here as a black dashed curve). Bars show daily disease incidence data, the solid line represents posterior mean estimates, and the shaded area represents central 95% credible intervals. B. Inferred time-dependent reproduction number, *R_t_*, for the suitability-based model in which *R_t_* = *C_t_S_t_* (estimates of the scaling factor, *C_t_*, are shown in Figure S2; blue), and for the autoregressive model with no explicit temperature-suitability component (orange). C. Daily risk of additional cases for the suitability-based (blue) and autoregressive (orange) models. Vertical dotted lines indicate the date on which emergency measures for blood collection were lifted (1 December [39]; red), and the date on which the 2025 ECDC criterion for a reduction in locally acquired *Aedes*-borne disease risk level would have been satisfied (20 December, 45 days after the last observed case occurred [8]; pink). The ECDC criterion was established eight years after the Anzio outbreak. D. Number of days from the last observed case to the risk of additional cases falling below a pre-specified threshold, shown for a range of threshold values.

Estimates of *R_t_* for the Anzio outbreak are shown in Figure 4B. We compared estimates obtained using our model accounting for seasonality (the suitability-based model; blue line and shaded region in Figure 4B) to comparable estimates obtained by fitting the renewal equation model to the data without explicit temperature dependence (the autoregressive model; orange), finding that the two models give very similar *R_t_* estimates during the acute phase of the outbreak (since substantial evidence about the value of *R_t_* is available from realised transmission reflected in the incidence data). However, *R_t_* estimates differ between the models from around the start of October: estimates for the autoregressive model begin to revert towards the prior distribution due to the lack of epidemiological data at the tail end of the outbreak, whereas estimates for the suitability-based model continue to decrease due to cooling temperatures.

We then calculated the daily risk of additional cases using the two fitted models (Figure 4C; each risk value incorporates the full posterior distribution for *R_t_* on and after the corresponding day).

Following the last observed case on 5 November, the suitability-based model indicates a substantially lower risk of additional cases than the autoregressive model, due to the seasonally forced decline in *R_t_* for the suitability-based model (Figure 4B). As a result, the suitability-based model indicates that interventions could theoretically have been relaxed earlier than suggested by the autoregressive model, based on the modelled risk of additional cases falling below a specified threshold (Figure 4D). For example, the risk of additional cases first falls below 1% after 19 days following the last observed case for the suitability-based model and 25 days for the autoregressive model.

In practice, the only explicit public indication of the relaxation of interventions in Anzio was the lifting of restrictions on blood donations on 1 December (26 days after the last observed case [39]). Both models suggest a very low risk of additional cases by the date on which restrictions on blood donations were lifted: 0.05% for the suitability-based model and 0.48% for the autoregressive model. By 45 days after the last observed case (the ECDC criterion for a reduction in *Aedes*-borne disease risk level, based on guidelines published years later in 2025 [8]), the models indicate zero risk of additional cases. This is because we assumed a maximum possible serial interval of 40 days (in fact, serial intervals of above 30 days are unlikely for chikungunya – see Figure 3D) and did not consider under-reporting in our initial analysis (this assumption is relaxed below).

In addition to comparing the suitability-based model with the autoregressive model (the equivalent model without explicit temperature dependence), we compared its estimates against two alternative reference approaches that do not include seasonality: (i) estimating *R_t_* using the autoregressive model, but then fixing posterior *R_t_* estimates from the date of the last observed case onwards at the estimate for that date (preventing estimates from reverting towards the prior distribution in the absence of new data; Figure S3A-C); and (ii) estimating *R_t_* using the EpiEstim R package [40–42] (Figure S3D-F). In both cases, the suitability-based model gives a lower risk of further cases after the last observed case than the reference model being considered, consistent with Figure 4.

We estimated the risk of additional cases in Figure 4 using *R_t_* estimates obtained by fitting the renewal equation to the entire disease incidence time series. However, we conducted an additional quasi real-time analysis, in which future *R_t_* values were estimated each day using only the disease incidence time series up to that day, and then used to calculate the risk of additional cases (Figure S4). While *R_t_* estimates exhibited greater day-to-day variations in the quasi real-time analysis (Figure S4C), since conducting a single fit to the entire incidence time series smooths out these variations, our finding that neglecting seasonality could lead to interventions being left in place longer than necessary was unchanged.

### Under-reporting extension

In our initial analysis of the Anzio outbreak (Figure 4), for simplicity and to isolate the effect of seasonality, we did not account for case under-reporting. We then relaxed this assumption, estimating under-reported case counts (shown for the suitability-based model in Figure 5A) alongside *R_t_* (Figure S5) under the assumption of a 60% reporting rate (consistent with estimates from a post-outbreak serosurvey of the 2007 Italian chikungunya outbreak [43]).

**Figure 5.**
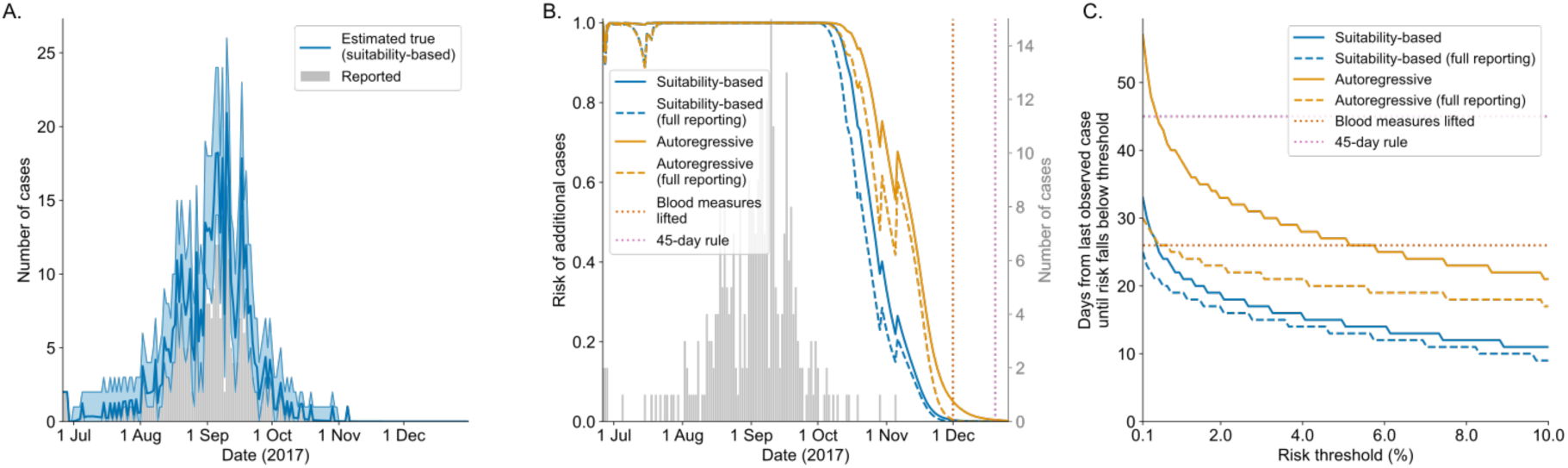
Application to the 2017 chikungunya outbreak in Anzio accounting for case under-reporting. A. Inferred disease incidence time series for the suitability-based model alongside reported data (bars), assuming a 60% reporting rate. The blue line represents posterior mean estimates, and the shaded area represents central 95% credible intervals. B. Daily risk of additional cases for the suitability-based (blue) and autoregressive (orange) models, assuming a reporting rate of either 60% (solid) or 100% (dashed). Vertical dotted lines indicate the date on which emergency measures for blood collection were lifted (1 December [39]; red), and the date on which the 2025 ECDC criterion for a reduction in locally acquired *Aedes*-borne disease risk level would have been satisfied (20 December, 45 days after the last observed case occurred [8]; pink). C. Number of days from the last observed case to the risk of additional cases falling below a pre-specified threshold, shown for a range of threshold values.

As would be expected, accounting for possible unreported cases led to higher estimates of the risk of additional cases under both the autoregressive and suitability-based models (Figure 5B). However, whereas accounting for under-reporting causes risk estimates for the autoregressive model to decline more slowly following the last reported case than when under-reporting is neglected, this effect is less pronounced for the suitability-based model. This is because accounting for the seasonal decline in transmission leads to lower posterior estimates of the number of unreported cases around and following the time of the last reported case. For example, risk estimates on 1 December (the date blood restrictions were lifted in Anzio) accounting for under-reporting are 0.40% for the suitability-based model (compared to 0.05% when assuming full reporting) and 5.06% for the autoregressive model (0.48% assuming full reporting). As a result, waiting times following the last reported case for the risk of additional cases to fall below a specified threshold differ more between the two models when accounting for under-reporting (Figure 5C). For example, for a 1% risk threshold, waiting times with under-reporting are 22 days for the suitability-based model (19 days assuming full reporting) and 39 days for the autoregressive model (25 days assuming full reporting). Consequently, accounting for under-reporting acted to enhance our main conclusion that failing to account for seasonality in transmission may lead to interventions being left in place longer than necessary at the end of an outbreak.

## Discussion

The decision to relax or remove interventions at the end of an infectious disease outbreak must balance social and economic costs of maintaining interventions against the risk of additional transmission. Here, we have analysed the role of seasonality in decision making at the end of an outbreak, showing that in scenarios with seasonal transmission, the time of year is an important determinant of when interventions can be relaxed. In such settings, a public health guideline that relaxes interventions a fixed period after the most recent case, while straightforward to implement and communicate, will not be appropriate at all times of the year: a fixed period adapted to the peak of the transmission season is likely to be unnecessarily long later in the season, whereas one suited to the end of the season may carry a high risk of additional cases if applied earlier. Our framework provides an alternative approach: using model estimates of the risk of further cases that account for seasonality and other factors such as case under-reporting, alongside the policymaker’s level of risk aversion, to guide decisions on when to relax interventions.

Following an *Aedes*-borne disease outbreak, ECDC guidance from 2025 recommends a 45-day waiting period from the symptom onset date of the last observed case before lowering the assigned risk level from level 3 (affected area) to level 2 (predisposed area) [8]. Once the risk level is reduced, the guidance indicates that surveillance and control measures, including active case finding, tracking of transmission chains, and vector control activities around cases, can be relaxed [8]. This 45-day period provides a risk-averse criterion under which the risk of additional cases after interventions are relaxed is low, at least in settings with effective clinical surveillance.

In operational settings, authorities could choose to relax some interventions earlier than the 45-day waiting period in order to balance the benefits of a conservative approach against the logistical, economic, and societal costs associated with prolonged control measures. This may be particularly beneficial around the end of the transmission season, since, as we have shown, long waiting periods at this time of year are often unnecessary. Following the 2017 chikungunya outbreak in Anzio, restrictions on blood donations were lifted on 1 December, 26 days after symptom onset in the last observed case [39], although we note that this was before the current ECDC guidance was published. Our suitability-based model, which explicitly accounts for seasonal variation in the transmission potential, estimates a very low risk of additional cases (0.40%) by the time that restrictions on blood donations were lifted during the Anzio outbreak, even under the assumption of substantial case under-reporting. These findings highlight how quantitative risk-based approaches may help support operational decision making by balancing precaution against the timely relaxation of interventions.

We acknowledge some limitations to our results. We assumed that *Aedes*-borne pathogen transmission could be modelled using a renewal equation based solely on human case data. We used seasonal temperature trends to inform estimates of *R_t_*, incorporating a lag to reflect that the number of human cases arising on a given day depends on environmental conditions over a human-vector-human transmission cycle ending on that day. However, our use of a smoothed seasonal temperature-suitability curve with a fixed lag is a simplification, since the number of cases arising on a given day is likely to depend on actual climate conditions over the entire transmission cycle [44] (although vectors may not respond immediately to small day-to-day temperature fluctuations). We also note that the temperature-suitability relationship [38] used as a prior in our suitability-based model is based on laboratory data collected under constant temperature, which may not fully capture real-world conditions or the ability of mosquitoes to exploit the anthropic environment. This may explain the cases occurring in Anzio in November 2017 despite a small seasonal suitability prior value. To address these limitations, we incorporated deviations from the seasonal trend in our approach, informed by observed transmission.

In summary, our modelling framework can be used to account for seasonality when estimating the risk of additional cases at the apparent end of an infectious disease outbreak. Future climate change, alongside other factors such as continuing urbanisation, is projected to drive further expansion of the spatial distribution of *Aedes* vectors [1,45], increasing the risk and geographic range of seasonal *Aedes*-borne disease outbreaks in temperate settings [7,46]. This highlights the importance of quantitative tools to support public health policymaking at all phases of *Aedes*-borne disease outbreaks, including towards their end. Incorporating seasonal transmission dynamics into end-of-outbreak risk assessment represents an important step towards strengthening evidence-based decision making for *Aedes*-borne diseases, as well as for other diseases with seasonal transmission dynamics.

## Methods

### Transmission model

We modelled transmission using a renewal equation [41,47,48],

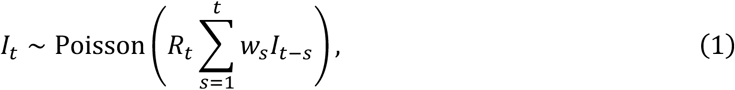

where *I_t_* is the incidence of new symptomatic human cases occurring on day *t* > 0 (with the number of index cases, *I*_0_, specified), *R_t_* is the time-dependent reproduction number, and *w_s_* is the probability that the serial interval takes the value *s* days. In the context of vector-borne disease, *R_t_* represents the expected total number of secondary human cases generated by each human case over the host-vector-host transmission cycle, assuming that future transmission conditions remain identical to those responsible for human cases arising on day *t*, and the values of *w_s_* similarly characterise the distribution of the serial interval over the host-vector-host cycle.

### Risk of additional cases

Given daily human disease incidence data up to day *t* (i.e., *I*_0_,…,*I_t_*_–1_), and assuming *R_t_* is known for all time, the risk of additional cases occurring on or after day *t* is given by

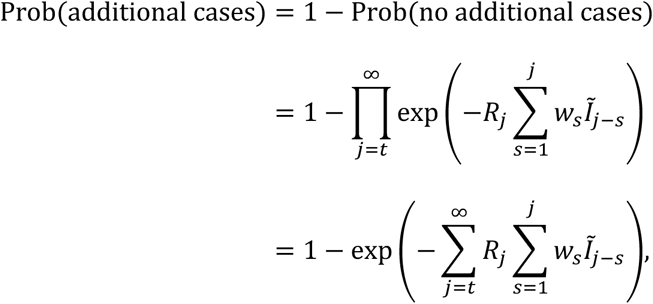

where *I*Fj–*s* is taken to be equal to *I*_j–*s*_ when 0 ≤ *j* − *s* < *t*, and zero otherwise (i.e., *I*Fj–*s* gives the complete disease incidence time series if no cases occur on or after day *t*). If the serial interval distribution has bounded support, i.e., *w_s_* = 0 for *s* > *s_max_*, then if *t*^∗^ ≤ *t* − 1 is the day of the most recent case, we have (for *t* ≤ *t*^∗^ + *s_max_*)

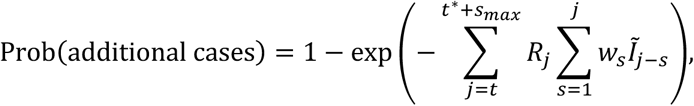

with zero risk of additional cases for *t* > *t*^∗^ + *s_max_*.

In a scenario where future values of *R_t_* are not known exactly, but *K* equally weighted posterior samples, denoted *R*_j_^(*k*)^ for *t* ≤ *j* ≤ *t*^∗^ + *s_max_* and 1 ≤ *k* ≤ *K*, are available (see “Anzio chikungunya outbreak case study” below), then the risk of additional cases accounting for uncertainty in *R_t_* is given by

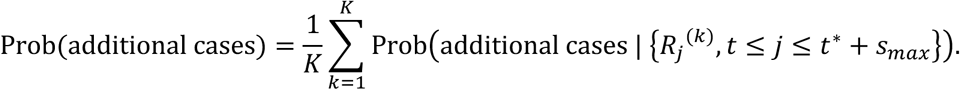

### Simulation study

In our initial analysis (Figure 2), we considered a scenario where *R_t_* = 2 × *S_seas_*(*d*_0_ + *t*), where *d*_0_ is the calendar day of the first case and *S_seas_* (which lies between 0 and 1) is the smooth seasonal suitability curve we estimated for chikungunya (see below; sensitivity to the precise seasonal transmissibility curve is considered in Figure S1). We also used the serial interval distribution for chikungunya described below. We first considered a single incident case, and then calculated on each subsequent day the risk of additional cases occurring on or after that day (assuming no cases occurred between the initial case and the current day, and assuming both the value of *R_t_* each day and the serial interval distribution to be known exactly). To verify the analytically derived risk values, we also estimated the risk of additional cases via renewal equation model simulations, using the incidence time series up to the day in question as the initial conditions for each simulation, and calculating the proportion of 10,000 simulations in which no cases occurred in the subsequent maximum serial interval, *s_max_*, days.

We then explored model-informed intervention relaxation decisions for a large number of synthetic outbreaks. Specifically, we generated each outbreak via simulation of the renewal equation model, starting with a single imported case occurring on a calendar day sampled uniformly at random over a full year, and continuing the simulation until the outbreak ended (i.e., an interval of *s_max_* days without cases had passed). If zero cases occurred after the index case (i.e., no locally infected cases), the simulation was discarded and the time of introduction resampled. We continued to conduct model simulations until 100,000 outbreaks with at least one locally infected case were obtained. For each synthetic outbreak, we calculated the number of days from the last observed case until the risk of additional cases fell below a 1% threshold.

### Anzio chikungunya outbreak case study

#### Outbreak data

We analysed disease incidence data from the chikungunya outbreak that occurred in Italy between June and November 2017 (grey bars in Figure 4A-C) [39]. Specifically, we used data from the primary outbreak cluster in the town of Anzio, comprising the symptom onset dates of 311 confirmed, probable and suspected cases (we excluded an additional reported case with unknown symptom onset date) [39]. We note that two cases were recorded on the first case day (26 June 2017); we treated these two cases as imported cases in our analyses, and assumed that subsequent cases arose as a result of local transmission.

#### Serial interval distribution

Using data from the 2017 Anzio chikungunya outbreak, Guzzetta *et al.* [37] estimated the continuous generation time distribution, describing intervals between infection times in human infector-infectee pairs (over the human-vector-human cycle), to follow a gamma distribution with shape parameter 8.53 and scale parameter 1.46 (mean 12.45 days, standard deviation 4.26 days). We assumed that the continuous serial interval (i.e., the interval between the precise symptom onset times of an infector and infectee) follows this same distribution, since Guzzetta *et al.* found that reconstructed serial intervals followed this distribution closely (see Figure S3 of [37]). We then discretised this distribution using the method described in [14,41], reassigning the probability of a zero-day serial interval (which is not permitted in the renewal equation model) to one day, and truncating the distribution at a maximum serial interval of *s_max_* = 40 days (reassigning the probability of a longer serial interval to this value). The discrete serial interval probability mass function is shown in Figure 3D.

#### Temperature suitability for transmission

We retrieved daily mean temperature data for 2010-2024 from the Rome Ciampino airport weather station using the Meteostat Python library [49]. We then obtained a smooth seasonal temperature curve by fitting a Fourier regression model with two harmonics,

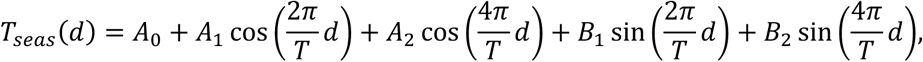

where *d* is the day of year and *T* = 365.25 days, to the data via least squares estimation using the statsmodels Python library [50] (Figure 3A).

We combined the seasonal temperature curve with the model describing temperature suitability for *Ae. albopictus*-borne chikungunya transmission (i.e., the relative reproduction number as a function of temperature, scaled to have a maximum value of one) by Tegar *et al.* [38] (Figure 3B). Specifically, Tegar *et al.* provide a grid of temperature-suitability combinations at a resolution of 0.1℃. Because *R_t_* in our renewal equation model corresponds to the square of that in [38] (since here, we consider the reproduction number for the full human-vector-human transmission cycle), we calculated the square of each grid suitability value, and then computed the seasonal suitability values, *S_seas_*(*d*), via linear interpolation (Figure 3C). Since the measured *R_t_* value on a given day is likely to reflect temperature conditions over the preceding host-vector-host transmission cycle, we computed *S_seas_*(*d*) based on the trend temperature value eight days previously, i.e., *T_seas_*(*d* − 8). This lag comprises the mean intrinsic (human) incubation period (estimated as three days [51]) plus half the average length of the remainder of the transmission cycle (the mean serial interval minus the mean intrinsic incubation period). Numerically, this is 3 + 0.5 × (12.45 − 3) days, rounded to the nearest integer.

#### Inference of the time-dependent reproduction number

We estimated *R_t_* by fitting the renewal equation model to the outbreak data from Anzio using Markov chain Monte Carlo, specifically the “nutpie” No-U-Turn Sampler (NUTS) provided by the PyMC Python library [52] (Figure 4A-B and Figure S2). We considered two model variants in our main analysis:

1. The autoregressive model, in which log(*R_t_*) = *ε_t_* is assumed a priori to follow an AR(1) process: *ε_t_*|*ε_t_*_–1_ ∼ Normal(*ρε_t_*_–1_, *σ*^2^(1 − *ρ*^2^)), with *ε*_0_ ∼ Normal(0, *σ*^2^), where we chose *σ* = 0.821 and *ρ* = 0.975. These hyperparameter choices ensure a stationary process, with the marginal prior distribution for *R_t_* on each day following a lognormal distribution with median one and 95% prior probability interval [0.2,5]. The relatively high value of *ρ* encodes prior belief that *R_t_* only varies slightly between successive days.
2. The suitability-based model, in which the time-dependent reproduction number is assumed to be of form *R_t_* = *C_t_S_t_*. Here, *S_t_* represents temperature suitability for transmission, assumed to be of form *S_t_* = min(max(*S_seas_*(*d*_0_ + *t*) + *ε_S_*_,*t*_, 0),1), where *d*_0_ is the day of year of the first case(s). The deviation from the seasonal suitability trend, *ε_S_*_,*t*_, was assumed a priori to follow an AR(1) process of the same form described above for the autoregressive model, except with hyperparameters *σ_S_* = 0.05 and *ρ_S_* = 0.975. This form for *S_t_* was chosen to restrict estimates to the range [0,1], while allowing a nonzero probability of either zero or maximum suitability. We note that in practice, we replaced the “hard clip” function min(max(⋅ ,0), 1) with a slightly smoothed “soft clip” function to ensure a differentiable likelihood (see the Supplementary Text). *C_t_* represents non-temperature factors affecting transmission, where log(*C_t_*) = *ε_C_*_,*t*_ was also assumed a priori to follow an AR(1) process of the same form, with hyperparameters *σ_C_* = 0.821 and *ρ_C_* = 0.975.

For each model, we ran four independent MCMC chains, each consisting of 1,000 tuning iterations (discarded as burn-in) followed by 1,000 retained iterations, resulting in a total of 4,000 equally weighted posterior samples of the *R_t_* time series. These settings yielded an effective sample size (ESS) of at least 1,000 for each *R_t_* estimate across time points and models in Figure 4, with all values of the *R*n statistic no greater than 1.01 (convergence diagnostics were computed using the ArviZ Python library [53]). We note that for the suitability-based model, *C_t_* and *S_t_* were estimated jointly, and posterior samples of *R_t_* were obtained as their product. We tested the inference procedure using simulated data (Figure S6).

In our analysis of the 2017 Anzio chikungunya outbreak presented in the main text, we fitted each model to the entire incidence time series. However, we also conducted an additional quasi real-time analysis in which the fitting process was repeated each day using only incidence data for earlier days (Figure S4). In this analysis, to calculate the risk of additional cases on/after a given day, *R_t_* estimates were obtained for the maximum serial interval, *s_max_* days, beyond that day.

### Under-reporting extension

To account for case under-reporting, we assumed that each incident case in the renewal equation (1) is reported with probability *q*. Writing the disease incidence as *I_t_* = *X_t_* + *Y_t_*, where *X_t_* and *Y_t_* denote reported and unreported cases developing symptoms on outbreak day *t*, respectively, by the Poisson thinning property we have

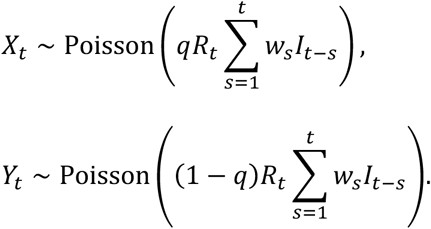

In our extended analysis of the Anzio chikungunya outbreak considering case under-reporting (Figure 5), we assumed a reporting probability of *q* = 0.6, chosen to be consistent with estimates from a post-outbreak serosurvey of the 2007 Italian chikungunya outbreak (in which 21 of 33 surveyed participants with chikungunya antibodies were reported during the outbreak) [43].

We fitted the extended renewal equation model to the disease incidence data, considering the same autoregressive and suitability-based model variants as described above, and estimating unreported case counts, *Y_t_*, alongside *R_t_*. For simplicity, given that our focus was on the latter stages of the outbreak, we assumed no unreported cases on or before the first reported case day (*t* = 0). Model fitting was implemented in PyMC, using a combination of NUTS (to update estimates of continuous-valued model parameters) and Metropolis (to update estimates of the discrete-valued unobserved case counts) steps. For each model, we ran four independent MCMC chains, each consisting of 2,000 tuning iterations followed by 8,000 retained iterations, which yielded ESS values of at least 1,000 and *̂R* values no greater than 1.01 for all estimated *R_t_* and *Y_t_* values.

In this analysis, the risk of additional cases was taken to be the probability of one or more cases (either reported or unreported) arising on or after the day of calculation, obtained by averaging the per-sample probability over all joint posterior samples of the *R_t_* and *Y_t_* time series.

## Data availability

All data are available at https://github.com/will-s-hart/end-of-outbreak-vbd (archived at https://doi.org/10.5281/zenodo.21260738). The epidemic curve for the Anzio chikungunya outbreak was previously published by Manica *et al.* [39].

## Code availability

All code used in the analyses is available at https://github.com/will-s-hart/end-of-outbreak-vbd (archived at https://doi.org/10.5281/zenodo.21260738). Computer code was written in Python (compatible with version 3.13).

## Acknowledgements

Thanks to members of the Wolfson Centre for Mathematical Biology at the University of Oxford, particularly the Infectious Disease Modelling group, for useful discussions about this work. This research was funded by Wellcome (grant number 226057/Z/22/Z; to W.S.H. and R.N.T.). R.N.T. is a member of the JUNIPER partnership, which is funded by MRC (grant number MR/X018598/1). The data collection and management were supported by funds allocated to the National Institute for Infectious Diseases “L. Spallanzani” (IRCCS), 00149 Rome (Italy) from the Italian Ministry of Health (Programme Ricerca Corrente – Linea 1 on emerging and re-emerging infections). For the purpose of Open Access, the authors have applied a CC BY public copyright licence to any Author Accepted Manuscript (AAM) version arising from this submission.

## Author contributions

W.S.H.: methodology, formal analysis, investigation, software, visualisation, writing (original draft), writing (review and editing).

C.M.: methodology, formal analysis, investigation, writing (review and editing).

M.M.: writing (review and editing).

F.M.: writing (review and editing).

M.S.: data collection, investigation, writing (review and editing).

F.V.: data collection, investigation, writing (review and editing).

P.P.: conceptualisation, writing (review and editing).

G.G.: conceptualisation, writing (review and editing).

R.N.T.: conceptualisation, methodology, supervision, writing (review and editing).

## Competing interests

The authors declare no competing interests.

## Notes

### Competing Interest Statement

The authors have declared no competing interest.

## References

1. Kraemer, M. U. G. et al. Past and future spread of the arbovirus vectors Aedes aegypti and Aedes albopictus. Nat. Microbiol. 4, 854–863 (2019).

2. Lim, A. et al. The overlapping global distribution of dengue, chikungunya, Zika and yellow fever. Nat. Commun. 16, 3418 (2025).

3. Caputo, B., et al. A comparative analysis of the 2007 and 2017 Italian chikungunya outbreaks and implication for public health response. PLoS Negl. Trop. Dis. 14, e0008159 (2020).

4. Buonfrate, D. et al. High burden of autochthonous arboviral infections during the summer season in Verona province, Italy, during 2025. J. Infect. 92, 106730 (2026).

5. European Centre for Disease Prevention and Control. Seasonal Surveillance of Chikungunya Virus Disease in the EU/EEA, Weekly Report. https://chik-weekly.ecdc.europa.eu/archive/chik-2025.html (2026).

6. Mordecai, E. A. et al. Thermal biology of mosquito-borne disease. Ecol. Lett. 22, 1690–1708 (2019).

7. Hart, W. S., et al. Climate variability amplifies the need for vector-borne disease outbreak preparedness. Proc. Natl. Acad. Sci. 122, e2507311122 (2025).

8. European Centre for Disease Prevention and Control. Public Health Guidance for Assessing and Mitigating the Risk of Locally-Acquired Aedes-Borne Viral Diseases in the EU/EEA. https://data.europa.eu/doi/10.2900/8091262 (2025).

9. Ministero della Salute. Piano Nazionale Di Prevenzione, Sorveglianza e Risposta Alle Arbovirosi (PNA) 2020-2025. https://www.salute.gov.it/new/it/pubblicazione/piano-nazionale-di-prevenzione-sorveglianza-e-risposta-alle-arbovirosi-pna-2020-2025/ (2019).

10 . Guzzetta, G., et al. Effectiveness and economic assessment of routine larviciding for prevention of chikungunya and dengue in temperate urban settings in Europe. PLoS Negl. Trop. Dis. 11, e0005918 (2017).

11. Djaafara, B. A. et al. A quantitative framework for defining the end of an infectious disease outbreak: application to Ebola virus disease. Am. J. Epidemiol. 190, 642–651 (2021).

12. World Health Organization. WHO Recommended Criteria for Declaring the End of the Ebola Virus Disease Outbreak. https://www.who.int/publications/m/item/who-recommended-criteria-for-declaring-the-end-of-the-ebola-virus-disease-outbreak (2020).

13. Hart, W. S. et al. Optimizing the timing of an end-of-outbreak declaration: Ebola virus disease in the Democratic Republic of the Congo. Sci. Adv. 10, eado7576 (2024).

14. Thompson, R. et al. Using real-time modelling to inform the 2017 Ebola outbreak response in DR Congo. Nat. Commun. 15, 5667 (2024).

15. Ogi-Gittins, I. et al. Real-time inference of the end of an outbreak: temporally aggregated disease incidence data and under-reporting. Infect. Dis. Model. 10, 935–945 (2025).

16. Akhmetzhanov, A. R., Jung, S., Cheng, H.-Y. & Thompson, R. N. A hospital-related outbreak of SARS-CoV-2 associated with variant Epsilon (B.1.429) in Taiwan: transmission potential and outbreak containment under intensified contact tracing, January–February 2021. Int. J. Infect. Dis. 110, 15–20 (2021).

17. Bradbury, N. V., Hart, W. S., Lovell-Read, F. A., Polonsky, J. A. & Thompson, R. N. Exact calculation of end-of-outbreak probabilities using contact tracing data. J. R. Soc. Interface 20, 20230374 (2023).

18. Eichner, M. & Dietz, K. Eradication of poliomyelitis: when can one be sure that polio virus transmission has been terminated? Am. J. Epidemiol. 143, 816–822 (1996).

19. Griette, Q., Liu, Z., Magal, P. & Thompson, R. N. Real-time prediction of the end of an epidemic wave: COVID-19 in China as a case-study. in Mathematics of Public Health: Proceedings of the Seminar on the Mathematical Modelling of COVID-19 (eds Murty, V. K. & Wu, J.) 173–195 (Springer International Publishing, Cham, 2022).

20. Lee, H. & Nishiura, H. Sexual transmission and the probability of an end of the Ebola virus disease epidemic. J. Theor. Biol. 471, 1–12 (2019).

21. Linton, N. M., Akhmetzhanov, A. R. & Nishiura, H. Localized end-of-outbreak determination for coronavirus disease 2019 (COVID-19): examples from clusters in Japan. Int. J. Infect. Dis. 105, 286–292 (2021).

22. Linton, N. M. et al. When do epidemics end? Scientific insights from mathematical modelling studies. Centaurus 64, 31–60 (2022).

23. Nishiura, H. Methods to determine the end of an infectious disease epidemic: a short review. in Mathematical and Statistical Modeling for Emerging and Re-emerging Infectious Diseases (eds Chowell, G. & Hyman, J. M.) 291–301 (Springer International Publishing, Cham, 2016).

24. Nishiura, H., Miyamatsu, Y. & Mizumoto, K. Objective Determination of End of MERS Outbreak, South Korea, 2015. Emerg. Infect. Dis. 22, (2016).

25. Parag, K. V. Sub-spreading events limit the reliable elimination of heterogeneous epidemics. J. R. Soc. Interface 18, 20210444 (2021).

26. Parag, K. V., Cowling, B. J. & Donnelly, C. A. Deciphering early-warning signals of SARS-CoV-2 elimination and resurgence from limited data at multiple scales. J. R. Soc. Interface 18, 20210569 (2021).

27. Parag, K. V., Donnelly, C. A., Jha, R. & Thompson, R. N. An exact method for quantifying the reliability of end-of-epidemic declarations in real time. PLOS Comput. Biol. 16, e1008478 (2020).

28. Plank, M. J. et al. Estimation of end-of-outbreak probabilities in the presence of delayed and incomplete case reporting. Proc. R. Soc. B Biol. Sci. 292, 20242825 (2025).

29. Thompson, R. N., Morgan, O. W. & Jalava, K. Rigorous surveillance is necessary for high confidence in end-of-outbreak declarations for Ebola and other infectious diseases. Philos. Trans. R. Soc. B Biol. Sci. 374, 20180431 (2019).

30. Yuan, B., Liu, R. & Tang, S. A quantitative method to project the probability of the end of an epidemic: Application to the COVID-19 outbreak in Wuhan, 2020. J. Theor. Biol. 545, 111149 (2022).

31. Carmona, P. & Gandon, S. Winter is coming: pathogen emergence in seasonal environments. PLOS Comput. Biol. 16, e1007954 (2020).

32. Hart, W. S. et al. Effects of individual variation and seasonal vaccination on disease risks. Nat. Commun. 16, 8471 (2025).

33. Kaye, A. R., Hart, W. S., Bromiley, J., Iwami, S. & Thompson, R. N. A direct comparison of methods for assessing the threat from emerging infectious diseases in seasonally varying environments. J. Theor. Biol. 548, 111195 (2022).

34. Kaye, A. R., Guzzetta, G., Tildesley, M. J. & Thompson, R. N. Quantifying infectious disease epidemic risks: a practical approach for seasonal pathogens. PLOS Comput. Biol. 21, e1012364 (2025).

35. Nadim, S. S., et al. Assessing risks of dengue, chikungunya and Zika transmission associated to Aedes albopictus in Chania, Greece, 2017–2018. PLoS Negl. Trop. Dis. 19, e0013785 (2025).

36. Vairo, F. et al. Local transmission of chikungunya in Rome and the Lazio region, Italy. PLOS ONE 13, e0208896 (2018).

37. Guzzetta, G. et al. Spatial modes for transmission of chikungunya virus during a large chikungunya outbreak in Italy: a modeling analysis. BMC Med. 18, 226 (2020).

38. Tegar, S., Brass, D. P., Purse, B. V., Cobbold, C. A. & White, S. M. Temperature-sensitive incubation, transmissibility and risk of Aedes albopictus-borne chikungunya virus in Europe. J. R. Soc. Interface 23, 20250707 (2026).

39. Manica, M., et al. Reporting delays of chikungunya cases during the 2017 outbreak in Lazio region, Italy. PLoS Negl. Trop. Dis. 17, e0011610 (2023).

40. Cori, A., et al. EpiEstim: Estimate Time Varying Reproduction Numbers from Epidemic Curves. (2026).

41. Cori, A., Ferguson, N. M., Fraser, C. & Cauchemez, S. A new framework and software to estimate time-varying reproduction numbers during epidemics. Am. J. Epidemiol. 178, 1505–1512 (2013).

42. Thompson, R. N. et al. Improved inference of time-varying reproduction numbers during infectious disease outbreaks. Epidemics 29, 100356 (2019).

43. Moro, M. L. et al. Chikungunya Virus in North-Eastern Italy: A Seroprevalence Survey. Am. J. Trop. Med. Hyg. 82, 508–511 (2010).

44. Mills, C. et al. Renewal equations for mosquito-borne diseases. Methods Ecol. Evol. 16, 2653– 2666 (2025).

45. Kaye, A. R. et al. The impact of natural climate variability on the global distribution of Aedes aegypti: a mathematical modelling study. *Lancet Planet*. Health 8, e1079–e1087 (2024).

46. Ryan, S. J., Carlson, C. J., Mordecai, E. A. & Johnson, L. R. Global expansion and redistribution of Aedes-borne virus transmission risk with climate change. PLoS Negl. Trop. Dis. 13, e0007213 (2019).

47. Fraser, C. Estimating individual and household reproduction numbers in an emerging epidemic. PLOS ONE 2, e758 (2007).

48. Ogi-Gittins, I. et al. A simulation-based approach for estimating the time-dependent reproduction number from temporally aggregated disease incidence time series data. Epidemics 47, 100773 (2024).

49. Lamprecht, C. S. Meteostat Python. https://dev.meteostat.net/python.

50. Seabold, S. & Perktold, J. Statsmodels: econometric and statistical modeling with Python. In Proceedings of the 9th Python in Science Conference 92–96 (2010).

51. Thiberville, S.-D. et al. Chikungunya fever: epidemiology, clinical syndrome, pathogenesis and therapy. Antiviral Res. 99, 345–370 (2013).

52. Abril-Pla, O. et al. PyMC: a modern, and comprehensive probabilistic programming framework in Python. PeerJ Comput. Sci. 9, e1516 (2023).

53. Martin, O. A. et al. ArviZ: a modular and flexible library for exploratory analysis of Bayesian models. J. Open Source Softw. 11, 9889 (2026).

