## Supplementary Material for "End-of-outbreak determination under seasonal transmission"

### Supplementary Material for "End-of-outbreak determination under seasonal transmission" by Hart *et al.*

#### Supplementary Text

<u>Soft clip function</u>

In our suitability-based model, temperature suitability for transmission is assumed to be of the form $S_t = \min(\max(\tilde{S}_t, 0), 1) = \mathrm{clip}(\tilde{S}_t; 0,1)$. Here, $\tilde{S}_t = S_{seas}(d_0 + t) + \varepsilon_{S,t}$, where $S_{seas}(d_0 + t)$ is the smoothed seasonal suitability, and $\varepsilon_{S,t}$ is an error term (assumed a priori to follow an autoregressive process). This construction was used to ensure a non-zero probability of suitability taking either its minimum or maximum values, but the non-differentiability of this hard clip function presents an issue for a No U-Turn Sampler (which requires a differentiable likelihood). Therefore, we replaced the hard clip with a smooth soft clip function as follows: the hard clip function can be written as

$$\mathrm{clip}(x; a, b) = \min(\max(x, a), b) = x + \max(a - x, 0) - \max(x - b, 0).$$

Now, a well-known smooth approximation to $\max(x, 0)$ is the softplus function:

$$\max(x, 0) \approx \mathrm{softplus}(x; \varepsilon) = \varepsilon \log(1 + e^{x/\varepsilon}),$$

where the approximation improves as the positive parameter, $\varepsilon$, becomes smaller. We therefore defined our soft clip function by

$$\mathrm{softclip}(x; a, b, \varepsilon) = x + \mathrm{softplus}(a - x; \varepsilon) - \mathrm{softplus}(x - b; \varepsilon).$$

We then took $S_t = \mathrm{softclip}(\tilde{S}_t; a = 10^{-8}, b = 1, \varepsilon = 0.001)$, where a lower bound slightly above zero was used to prevent suitability values reaching zero at machine precision, which results in a zero likelihood (and thus log-likelihood of $-\infty$).

#### Supplementary Figures

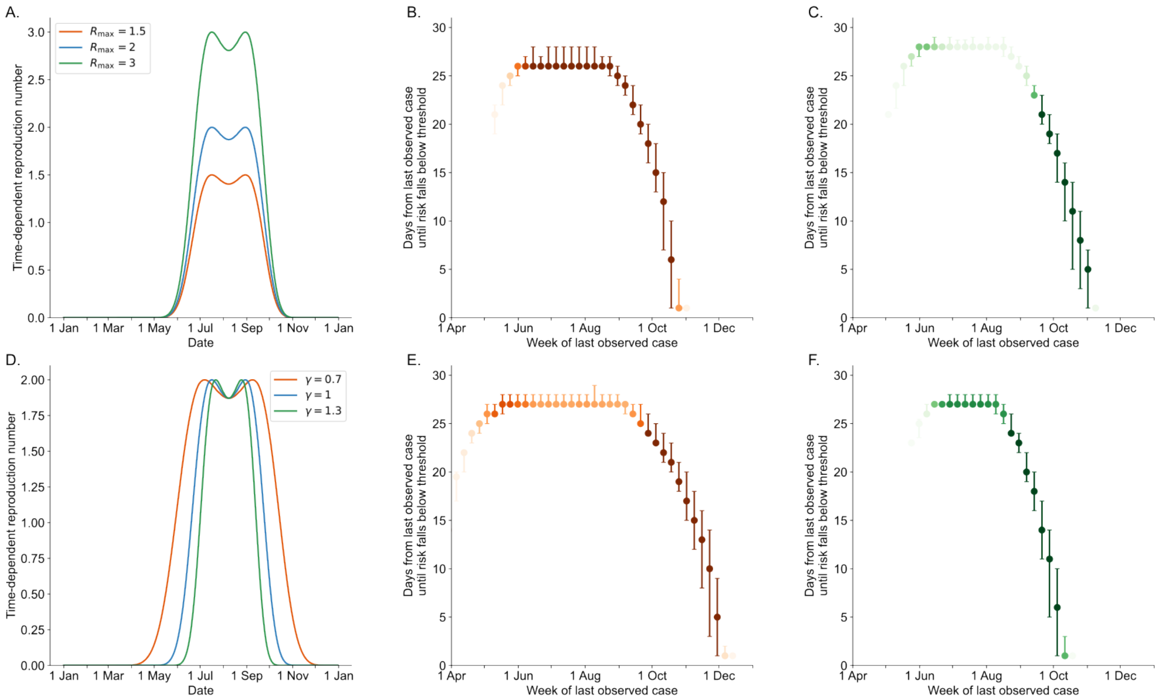


**Figure S1. Sensitivity of simulation study results to seasonal transmissibility profile.** We assumed a time-dependent reproduction number on outbreak day $t$ (calendar day $d_0 + t$) of form $R_t = R_{max} \times S_{seas}\left(d^* + \gamma \times (d_0 + t - d^*)\right)$, where $S_{seas}$ is the default seasonal suitability curve used in Figure 2, $R_{max}$ is the maximum possible reproduction number (default value 2), $d^* = 31$ July represents the middle of the transmission season (between twin summer peaks) and $\gamma$ is a parameter characterising the rate at which transmission declines away from its summer peaks (so that higher values of $\gamma$ correspond to a shorter transmission season; default value 1). Note that $S_{seas}(x)$ was computed for non-integer $x$ by linear interpolation, and was defined to be zero more than six months from $d^*$ (rather than continuing to wrap periodically) to ensure a single annual transmission season when $\gamma > 1$. A. Seasonal profile of $R_t$ for $R_{max}$ values of 1.5 (red), 2 (blue) and 3 (green), with $\gamma = 1$. B-C. Number of days from the last observed case until the risk of additional cases falls below 1%, shown by week of last observed case across 100,000 simulated outbreaks, as in Figure 2D, for $R_{max} = 1.5$ (B) and $R_{max} = 3$ (C), with $\gamma = 1$. D. Seasonal profile of $R_t$ for $\gamma$ values of 0.7 (red), 1 (blue) and 1.3 (green), with $R_{max} = 2$. E-F. Equivalent panels to B-C, but with $\gamma = 0.7$ (E) and $\gamma = 1.3$ (F), with $R_{max} = 2$.

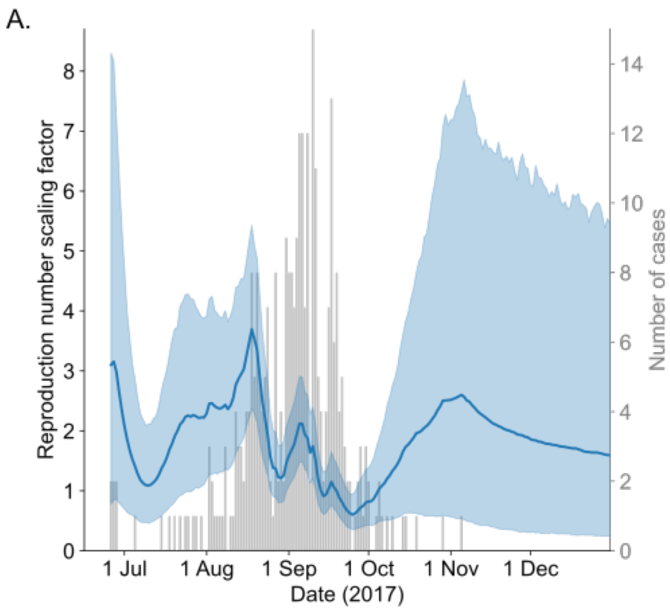


**Figure S2. Inferred reproduction number scaling factor for the 2017 Anzio chikungunya outbreak.** The reproduction number scaling factor, $C_t$, for the suitability-based model, was assumed a priori to follow an AR(1) process on the log scale. Bars show daily disease incidence data, the solid line represents posterior mean estimates, and the shaded area represents central 95% credible intervals.

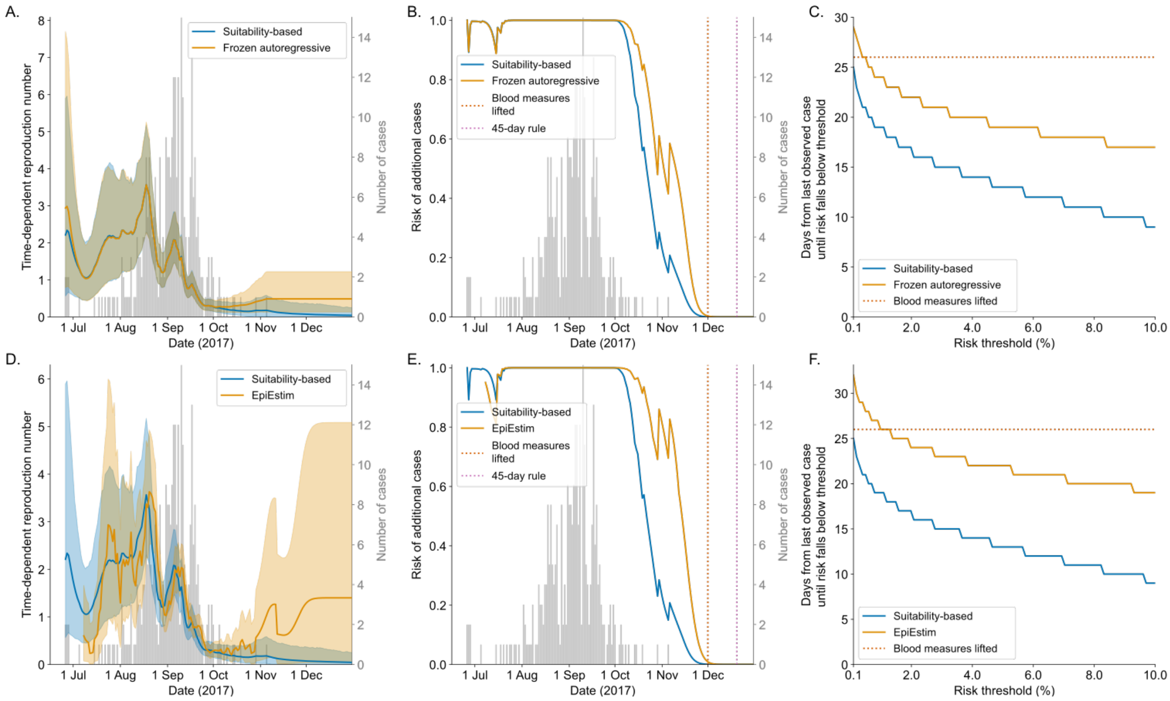


**Figure S3. Inference and end-of-outbreak determination for the 2017 Anzio chikungunya outbreak with alternative reference models.** A. Reproduction number, $R_t$, for the suitability-based model (blue) and for the autoregressive model but with posterior estimates following the date of the final observed case frozen at the estimate for that date (orange). B. Daily probability of additional cases. Vertical dotted lines indicate the date on which emergency measures for blood collection were lifted (1 December; red), and the date on which the 2025 ECDC criterion for a reduction in locally acquired *Aedes*-borne disease risk level would have been satisfied (20 December, 45 days after the final case occurred; pink). C. Number of days from the last observed case to the risk of additional cases falling below a pre-specified threshold, shown for a range of threshold values. D-F. Equivalent panels to A-C, but comparing the suitability-based model with results based on $R_t$ estimates obtained using the EpiEstim R package. To estimate $R_t$ with EpiEstim, we assumed a gamma prior with mean 1.40 and standard deviation 1.37 (matching the marginal $R_t$ priors in our autoregressive model), and a seven-day sliding window (i.e., each daily estimate assumes constant $R_t$ over that day and the preceding six; the sharp drop in the $R_t$ estimate a week after the last observed case reflects that case leaving the window).

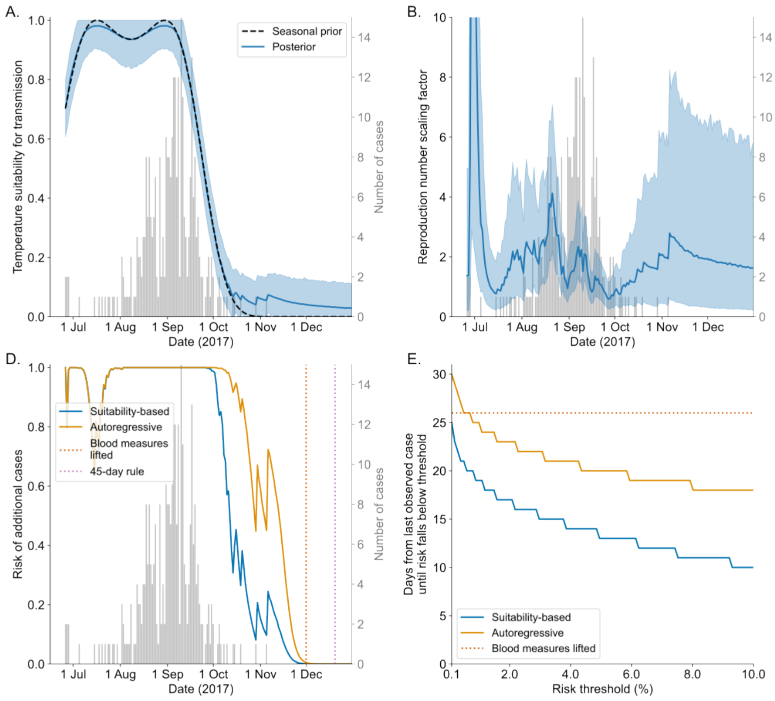


**Figure S4. Quasi real-time inference and end-of-outbreak determination for the 2017 Anzio chikungunya outbreak.** A. Temperature suitability for transmission, $S_t$. Bars show daily disease incidence data, the solid line represents posterior mean estimates, and the shaded area represents central 95% credible intervals. The black dashed curve shows the seasonal suitability profile used to inform model estimates. B. Reproduction number scaling factor, $C_t$, for the suitability-based model. C. Reproduction number, $R_t$, for the suitability-based (blue) and autoregressive (orange) models. D. Daily probability of additional cases. Vertical dotted lines indicate the date on which emergency measures for blood collection were lifted (1 December; red), and the date on which the 2025 ECDC criterion for a reduction in locally acquired *Aedes*-borne disease risk level would have been satisfied (20 December, 45 days after the final case occurred; pink). E. Number of days from the last observed case to the risk of additional cases falling below a pre-specified threshold, shown for a range of threshold values**.**

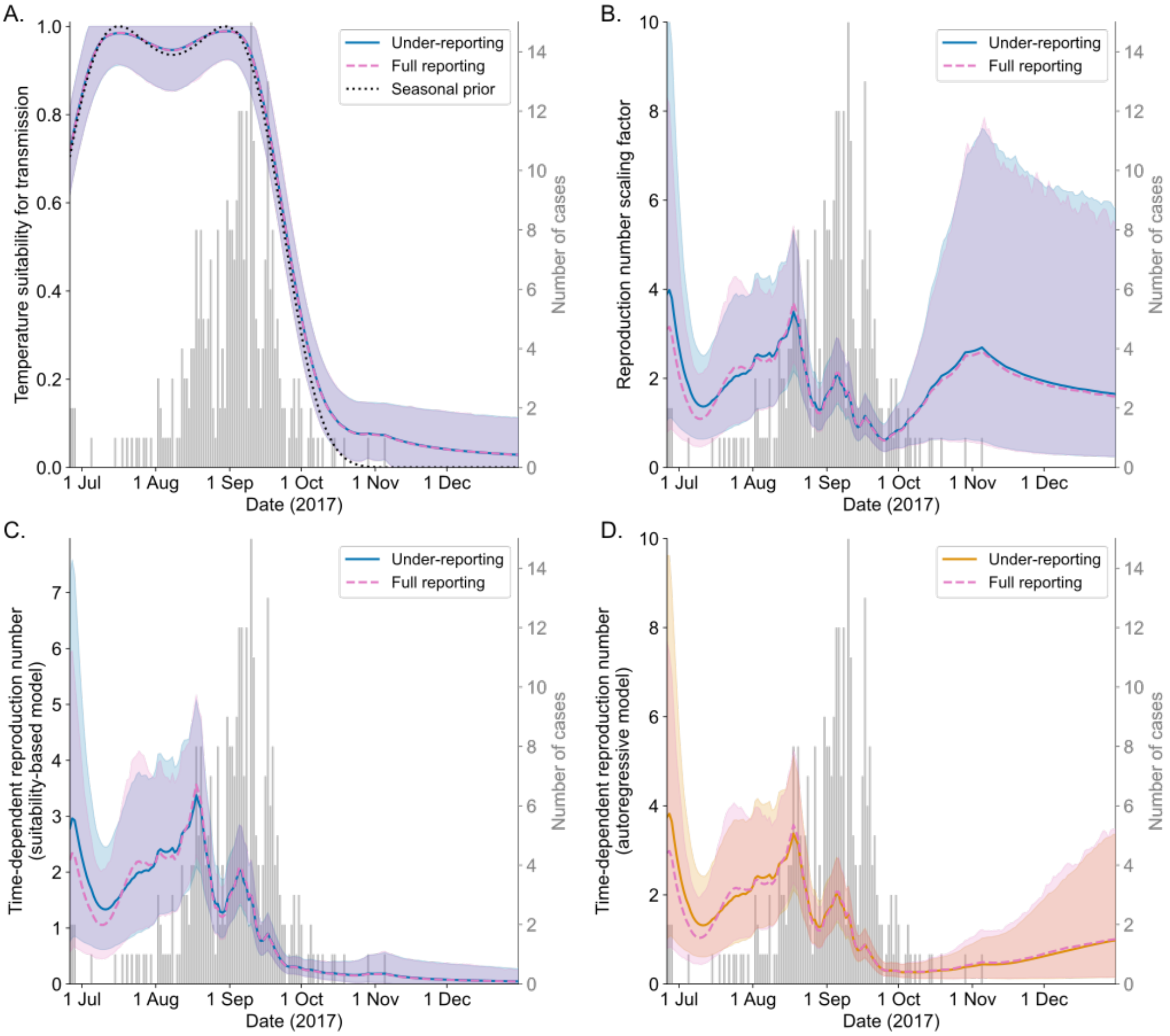


**Figure S5. Reproduction number inference for the 2017 Anzio chikungunya outbreak accounting for case under-reporting.** A. Temperature suitability for transmission, $S_t$. Bars show daily reported disease incidence data, lines represent posterior mean estimates assuming a reporting rate of either 60% (blue solid) or 100% (pink dashed), and shaded areas represent central 95% credible intervals. The black dotted curve shows the seasonal suitability profile used to inform model estimates. B. Reproduction number scaling factor, $C_t$, for the suitability-based model. C. Reproduction number, $R_t = C_t S_t$, for the suitability-based model. D. Reproduction number, $R_t$, for the autoregressive model (posterior mean estimates accounting for under-reporting are here shown in orange).

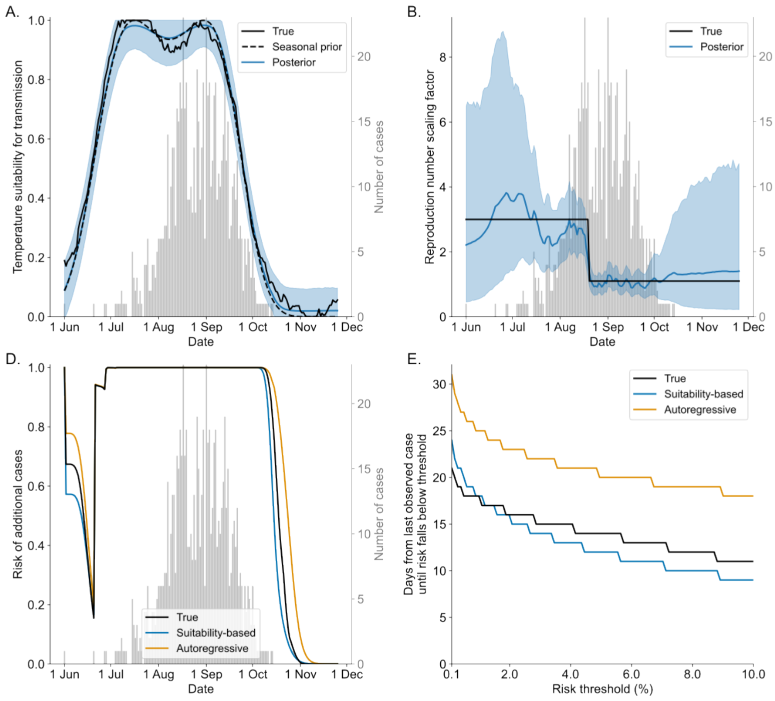


**Figure S6. Reproduction number inference and end-of-outbreak determination for a simulated outbreak**. A. Temperature suitability for transmission, $S_t$. Bars show simulated incidence data, the blue line shows posterior mean estimates, and the shaded area represents central 95% credible intervals. True values (solid black curve) were obtained by simulating the autoregressive process used as a prior for estimation of $S_t$ in our main analysis (see Methods). The black dashed curve shows the assumed seasonal suitability profile on which true and estimated suitability values were based. B. Reproduction number scaling factor, $C_t$, for the suitability-based model. True values were assumed to follow a step function (as shown). C. Reproduction number, $R_t$, for the suitability-based (blue) and autoregressive (orange) models. True values correspond to the product of the true values of $S_t$ and $C_t$. D. Daily probability of additional cases. E. Number of days from the last observed case to the risk of additional cases falling below a pre-specified threshold, shown for a range of threshold values**.**
